# Rejuveinix Reverses Severe Inflammatory Lung Damage in a Mouse Model of Fatal Sepsis

**DOI:** 10.1101/2020.12.31.424986

**Authors:** Fatih M. Uckun, Ibrahim H. Ozercan, Cemal Orhan, Marcus Gitterle, Kazim Sahin

**Affiliations:** Drug Discovery Program, Reven Pharmaceuticals, Westminster, CO 80234, USA; Department of Developmental Therapeutics, Immunology, and Integrative Medicine, Ares Pharmaceuticals, St. Paul, MN 55110, USA; Department of Pathology Faculty of Medicine, Firat University, Elazig 23119, Turkey; Department of Animal Nutrition, Faculty of Veterinary, Firat University, Elazig 23119, Turkey; and; Department of Surgery, Christus Health System, Santa Rosa Hospital, New Braunfels, TX 78130, USA

**Keywords:** Cancer, Sepsis, Pneumonia, ARDS, Acute lung injury (ALI), Multi-organ dysfunction (MODS), Cytokine release syndrome (CRS), COVID-19

## Abstract

We set out to determine if our anti-sepsis drug candidate Rejuveinix (RJX) can effectively treat systemic inflammation in a mouse model of sepsis when used at a dose level that is >10-times lower than its maximum tolerated dose (MTD) for human subjects. Our findings obtained in the LPS-GalN mouse model of fatal sepsis provide experimental evidence that RJX exhibits potent single-agent anti-inflammatory activity and reverses very severe inflammatory lung injury when used therapeutically, thereby significantly improving the survival outcome. These findings demonstrate the clinical impact potential of RJX for the treatment of severe sepsis.

## Introduction

Severe viral sepsis caused by SARS-CoV-2, the causative agent of coronavirus disease 2019 (COVID-19), shows a rapid progression associated with a cytokine release syndrome (CRS) and a high case fatality rate due to the development of ARDS and multi-organ failure in high-risk COVID-19 patients (1,2). Our anti-sepsis drug candidate, Rejuveinix (RJX), is a rationally-designed formulation of naturally occurring antioxidants and anti-inflammatory compounds with a significant clinical impact potential for COVID-19 associated viral sepsis (3-5). RJX showed a very favorable clinical safety profile in a recently completed double-blind, placebo-controlled, randomized, two-part, ascending dose-escalation Phase 1 study in healthy volunteers (Protocol No. RPI003; ClinicalTrials.gov Identifier: NCT03680105) (5). It has now entered a randomized double-blind, placebo-controlled Phase 2 study in hospitalized patients with critical COVID-19.

The goal of this research project was to determine if RJX can effectively treat systemic inflammation in a mouse model of sepsis. Our findings obtained in the LPS-GalN mouse model of fatal sepsis provide experimental evidence that RJX exhibits potent single-agent antiinflammatory activity and reverses very severe inflammatory lung injury when used therapeutically, thereby significantly improving the survival outcome.

## Results

LPS-GalN challenged mice experience a very rapidly progressing systemic inflammation with markedly elevated inflammatory cytokine levels as well as severe oxidative stress and acute lung injury at 2 hours post LPS-GalN injection. Histopathological examination of the lung tissues from LPS-GalN injected mice showed histological changes consistent with severe acute ALI, including alveolar hemorrhage, thickening of alveolar wall, edema/congestion, and leukocyte infiltration. The use of low dose RJX (Dose level: 0.7 mL/kg; Human equivalent dose = 0.057 mL/kg) corresponding to 7.5% of its clinical maximum tolerated dose (MTD) of 0.759 mL/kg (5) as a prophylactic anti-sepsis regimen was highly effective in preventing the development of very severe lung damage after injection with an invariably fatal dose of LPS-GalN (**Figure 1**).

**Figure 1.**
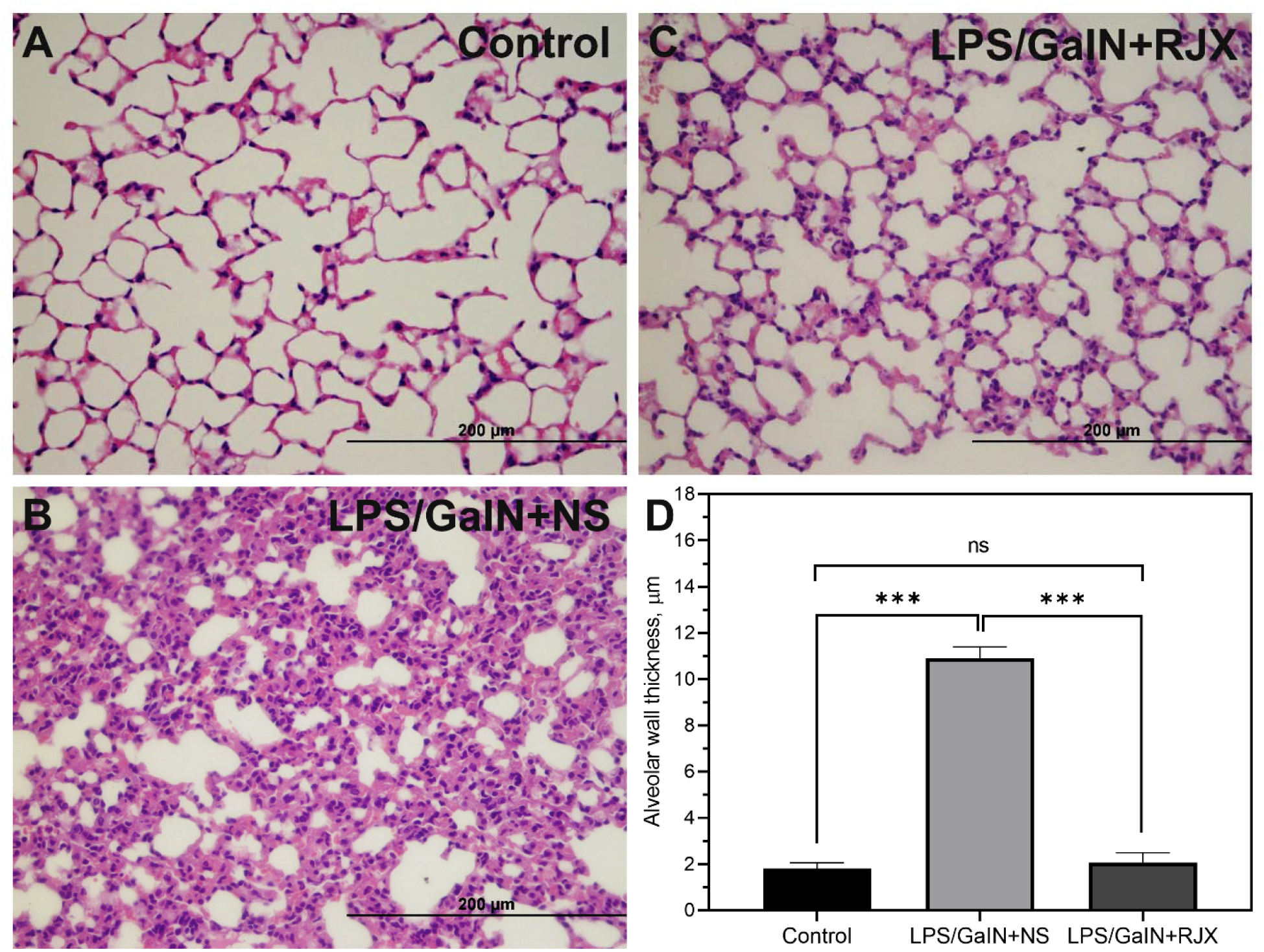
Prophylactic RJX Prevents Acute Lung Injury, Inflammation, and Pulmonary Edema in the LPS-GalN Mouse Model of ARDS and Multi-organ Failure. Mice were treated with i.p injections of 6-fold diluted RJX (4.2 mL/kg, 0.5 ml/mouse) (Panel C) or vehicle (NS) (Panel B) 2 hours before and 2 hours post-injection of LPS-GalN. Except for untreated mice (Control) (Panel A), each mouse received 0.5 ml of LPS-GalN (consisting of 100 ng of LPS plus 8 mg of D-galactosamine) i.p. The RJX dose levels (in mL/kg) are indicated in parentheses. The average thickness of the alveolar wall, depicted in a bar graph representation of the mean ±SD values, was determined as a measure of pulmonary edema (Panel D). H&E X400

Likewise, the therapeutic use of 0.7 mL/kg RJX after the onset of very severe lung damage and fulminant cytokine storm was highly effective in reversing the inflammatory damage in the lungs. Delayed treatments with RJX starting at 2 hours after the LPS-GalN injection attenuated the LPS-GalN induced acute lung injury and markedly reduced the pulmonary edema (**Figure 2**).

**Figure 2.**
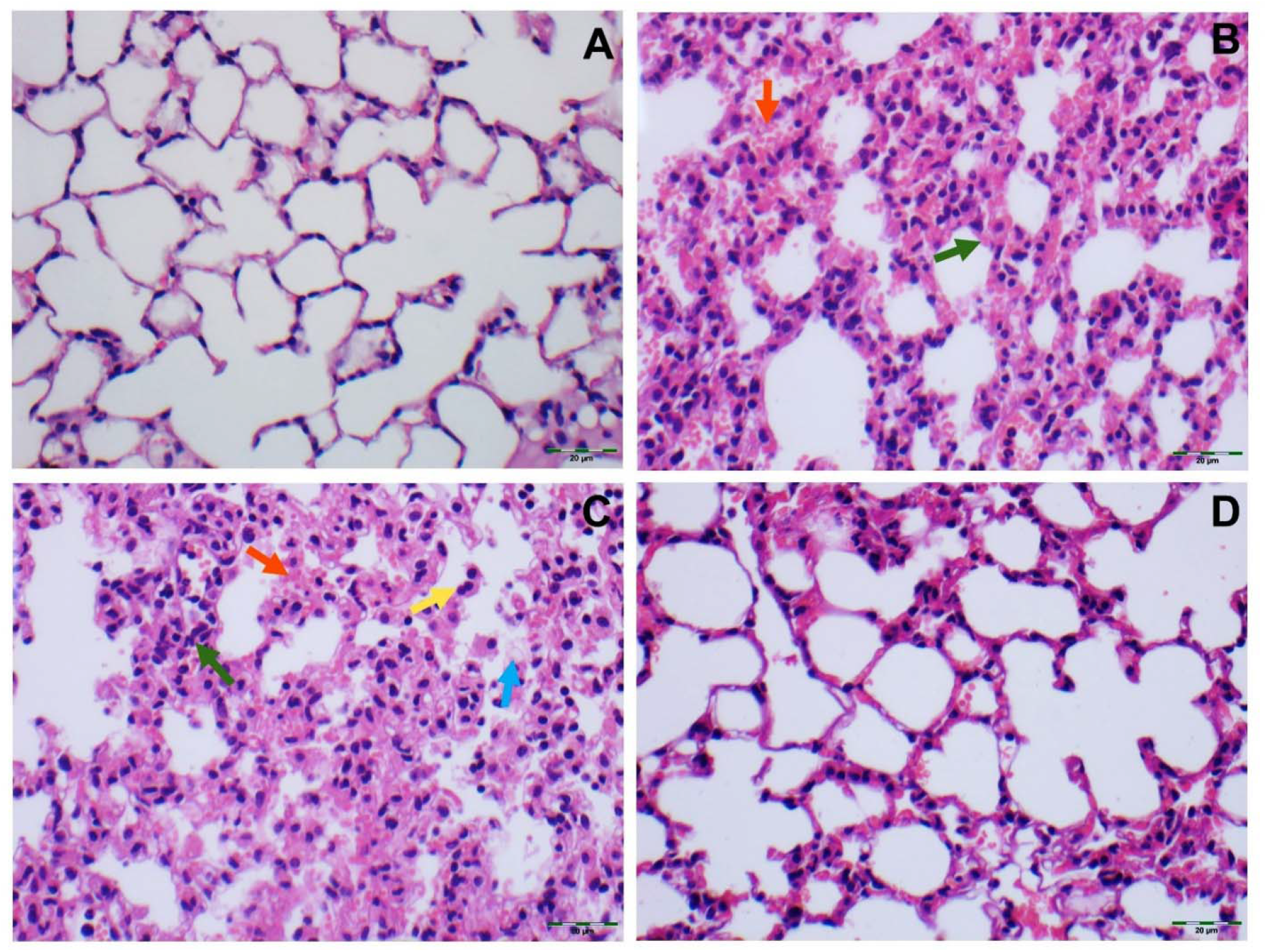
Therapeutic RJX Reverses Acute Lung Injury and Inflammation in a Mouse Model of Fatal Cytokine Storm, Sepsis, Systemic Inflammation, ARDS, and Multiorgan Failure. Mice were treated with i.p injections of RJX (6-fold diluted, 4.2 mL/kg, 0.5 ml/mouse) or NS (0.5 ml/mouse) 2 and 3 hours post-injection of LPS-GalN. Except for untreated control mice (Control, A, term at 2h), each mouse received 0.5 ml of LPS-GalN (consisting of 100 ng of LPS plus 8 mg of D-galactosamine) i.p. Groups of 6 BALB/C mice were treated with i.p injections of RJX (D, term at 24h), RJX or vehicle (NS) 2 hours post-LPS-GalN (1st treatment, B, term at 2h) and 3 hours post-LPS-GalN (2nd treatment, C, term at 24h). Yellow arrow: inflammatory cell infiltration; Blue arrow: Exudate; Orange arrow: hemorrhage; Green block: the thickness of alveolar wall. H&E X400

Treatment with RJX significantly improved the survival outcome. Control mice (n=6) treated with 2 injections of placebo (normal saline) at 2 hours and 3 hours, respectively, post-LPS-GalN challenge all rapidly died within 4 hours at a median survival time of 2.2 hours after the first injection of normal saline and 4.2 hours after the LPS-GalN challenge (**Figure 3**). Notably, treatment of mice with 0.7 mL/kg RJX at 2 hours and 3 hours post LPS-GalN injection resulted in improved survival outcome with 3 of 6 mice remaining alive at 24 hours post LPS-GalN injection (**Figure 3**). The median survival time of these mice was markedly longer than the median survival of normal saline-treated control mice (15.1 hours vs. 4.2 hours, P = 0.0098) (**Figure 3**).

**Figure 3.**
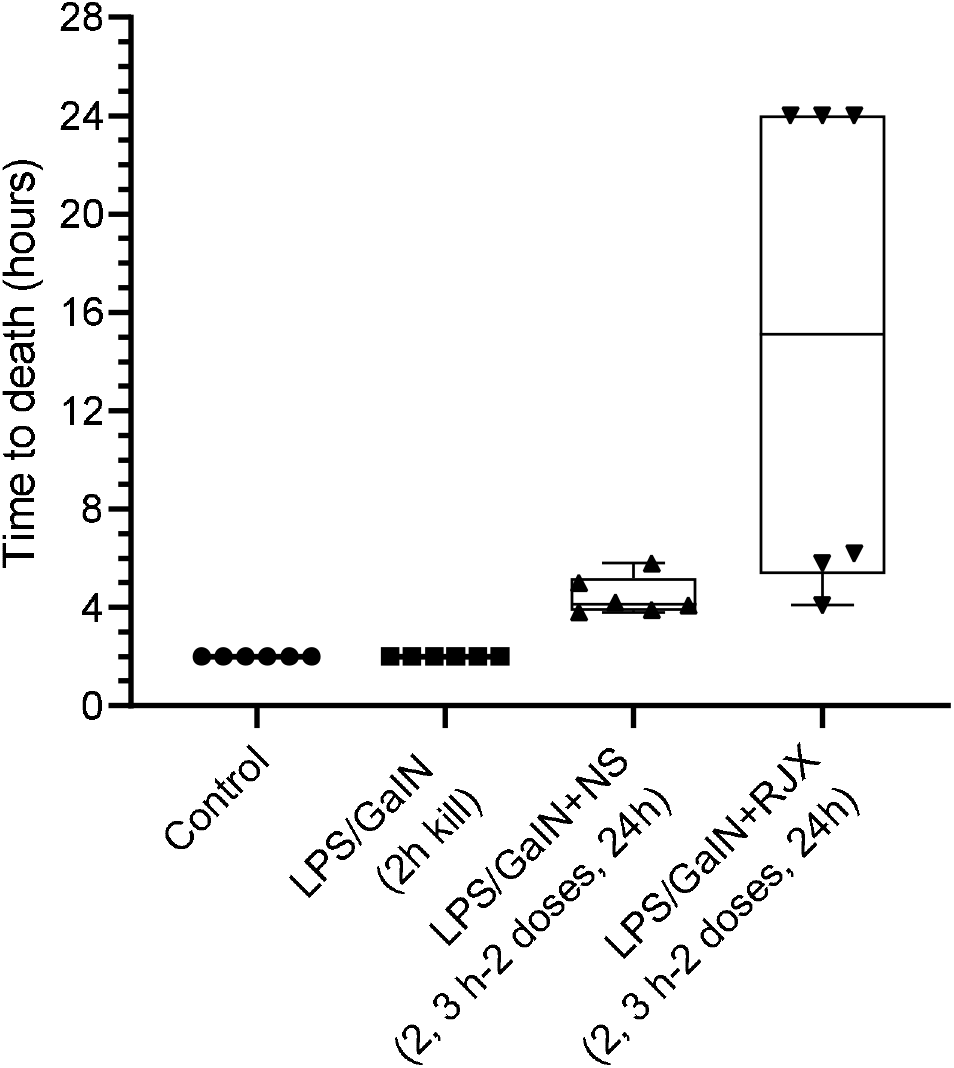
In Vivo Protective Activity of Delayed-Onset RJX Treatments in the LPS-GalN Model of ARDS and Multiorgan Failure. Groups of 6 BALB/C mice were treated with i.p injections of 6-fold diluted RJX (4.2 mL/kg, 0.5 ml/mouse) or vehicle (NS) 2 and 3 hours post-injection of LPS-GalN. Each mouse received 0.5 ml of LPS-GalN (consisting of 100 ng of LPS plus 8 mg of D-galactosamine) i.p. Percent (%) survival for each treatment group is shown as a function of time after the LPS-GalN challenge. The depicted Whisker plots represent the median and values for time to death from 6 mice in each group.

## Discussion

Here we report preclinical proof of concept data regarding the clinical impact potential of RJX for the treatment of sepsis. Specifically, we demonstrate that our anti-sepsis drug candidate RJX – at a dose level >10-fold lower than its clinical MTD-exhibits potent single-agent anti-inflammatory activity in the LPS-GalN mouse model of sepsis and multi-organ failure. It prevented inflammatory lung injury when used prophylactically and it reversed lung injury when used therapeutically, thereby significantly improving the survival outcome. The clinical potential of RJX is currently being examined in hospitalized COVID-19 patients with viral sepsis to test the hypothesis that will contribute to a faster resolution of respiratory failure and a reduced case mortality rate. A Phase II proof of concept study has also been designed to determine if RJX can favorably alter the clinical course and reduce the mortality rate of viral sepsis in critically ill COVID-19 patients.

## Methods

### Rejuveinix (RJX)

The ingredients of RJX include ascorbic acid, magnesium sulfate heptahydrate, cyanocobalamin, thiamine hydrochloride, riboflavin 5’ phosphate, niacinamide, pyridoxine hydrochloride, and calcium D-pantothenate. RJX is a two-vial system and A and B are each of the two vials. Vial A contains the active ingredients and minerals whereas Vial B contains the buffer, sodium bicarbonate as the Vial A content is acidic (5).

### LPS-GalN Model of Fatal Cytokine Storm, Sepsis, and Multi-organ Failure

The ability of RJX to prevent fatal shock, ARDS, and multi-organ failure was examined in the well-established LPS-GalN model (5). The animal research study was approved by the Animal Care and Use Committee of Firat University. BALB/c mice were randomly divided into different treatment groups using pseudo-randomization convenience allocation and we applied the concealment of treatment allocation and blind outcome assessment to reduce the risk of bias in our conclusions. Each mouse except for control mice received a 500 μL i.p. injection of LPS-GalN (consisting of 100 ng of LPS plus 8 mg of D-galactosamine). In the therapeutic setting, vehicle control mice were treated with 0.5 mL normal saline (NS), i.e., an aqueous solution of 0.9% NaCl instead of RJX intraperitoneally (*i.p*) 2 hours and 3 hours after the *i.p* injection of LPS-GalN. Test mice received 0.7 mL/kg RJX dose (2 hours and 3 hours after LPS-GalN. In the prophylactic setting, RJX was administered 2 hours before and 2 hours after the injection of LPS-GalN, as reported (5). Mice were monitored for mortality for 24 h. The Kaplan-Meier method, log-rank chi-square test, was used to analyze the 24 h survival outcomes of mice in the different treatment groups. At the time of death, lungs were harvested, fixed in 10% buffered formalin, and processed for histopathologic examination. 3 μm sections were cut, deparaffinized, dehydrated, and stained with hematoxylin and eosin (H & E), and examined with light microscopy.

### Statistical Analyses

Statistical analyses employed standard methods, including analysis of variance (ANOVA) and/or, nonparametric analysis of variance (Kruskal-Wallis) using the SPSS statistical program (IBM, SPPS Version 21), as reported (5). Furthermore, the Kaplan-Meier method, log-rank chi-square test, was used to investigate survival and fatality in each group.

## Author contributions

Each author has made significant contributions to the study, reviewed and revised the manuscript, approved the final version for submission. F.M.U conceived the study, designed the evaluations reported in this paper, directed the data compilation and analysis, analyzed the data, and prepared the initial draft of the manuscript. Each author had access to the source data used in the analyses I.H.O. performed the necropsies and histopathologic examinations on mice.

## Competing interests

F.M.U. is an employee of Reven Pharmaceuticals, the sponsor for the clinical development of RJX. I.H.O, C.O., M.G., and K.S declare no current competing financial interests.

